# The ocellate river stingray (*Potamotrygon motoro*) exploits vortices of sediment to bury into the substrate

**DOI:** 10.1101/834986

**Authors:** S.G. Seamone, D.A. Syme

## Abstract

Particle image velocimetry and video analysis were employed to discern and describe the mechanism used by the stingray *Potamotrygon motoro* to bury into the substrate. *P. motoro* repeatedly and rapidly pumped the body up and down while folding the posterior portion of the pectoral fins up and over, drawing water in and suspending sediment beneath the pectoral disc. As the fins folded up and over, vortices of fluidized sediment travelled along the ventral surface of the fins toward the fin tips, and were then directed onto the dorsal surface of the fins and towards the dorsal midline of the fish, where they dissipated and the sediment settled over the dorsal surface of the ray. As displacement and speed of the body pumping and finbeat motions increased, the speed of the sediment translating across the dorsal surface increased, and accordingly, sediment coverage of the dorsal surface increased. Mean sediment coverage was 82.5% ± 3.0 S.E.M, and appeared to be selectively controlled, whereby the pectoral fins tended to bury more than the body, head and tail, and the body more than the head and the tail. In the most vigorous burying events, vortices of sediment shed from each fin collided at the midline and annihilated, reorienting the sediment flow and sending jets of sediment towards the head and the tail, covering these locations with sediment. Hence, this study demonstrates that the mechanism of burying employed by *P. motoro* permits effective control of sediment vortices and flows to modulate the extent of burying.

## INTRODUCTION

Exploring burying behaviours in fishes provides insight into the relationships between shape, function, sediment dynamics and performance used for crypsis and station holding in a benthic, aquatic environment. Although our understanding of the diversity in the mechanisms used by fishes to bury and cover themselves with substrate is incomplete, the mechanics involved in burying appear to be closely related to their swimming mechanics. Fishes that use lateral undulations of the body to swim employ similar side-to-side bending movements of the body and caudal fin (i.e. BCF undulations) to displace sediment and bury into the substrate, often penetrating the substrate with either the head (Gidmark et al., 2011; Tatom-Naecker and Westneat, 2018) or ventral surface. Flatfish (i.e. pleuronectiformes), which are laterally flattened and flipped on their side, also swim via BCF undulations, but the undulations are oriented vertical relative to the substrate (Webb, 2002), and the vertical waves of bending fluidize the sediment underneath the animal, moving it into suspension and up and over onto the upper surface of the fish (Corn et al., 2018; Gibson and Robb, 1992; McKee et al., 2016; Stoner and Ottmar, 2003; Tanda, 1990). Stingrays (Suborder Myliobatoidei) are another group of fishes that are flattened in the same plane as the substrate and are common to benthic habitats in both freshwater and marine environments (Allen and Pondella, 2006; Aschliman et al., 2012; Goes de Araújo et al., 2004; Vaudo and Lowe, 2006), and are known to bury into the substrate. However, unlike flatfish which are flattened and flexible in the lateral axis, benthic stingrays are dorsoventrally flattened with a stiffened body axis and enlarged and rounded pectoral fins that wrap around the head and body forming the pectoral disc (Blevins and Lauder, 2012; Fontanella et al., 2013; Franklin et al., 2014; Parson et al., 2011; Schaefer and Summers, 2005). Furthermore, rather than axial bending, stingrays employ flexible movements of the pectoral fins to power routine and fast-start swimming (Blevins and Lauder, 2012; Rosenberger, 2001; Rosenberger and Westneat, 1999; Seamone et al., 2019). Hence, it might be anticipated that mechanisms of burying in stingrays will differ from flatfish and BCF swimming fishes, and exploring burying behaviours in stingrays might reveal an alternative approach that is used by fish to displace and settle sediment over a flattened and high surface area.

The aim of this study was to describe the mechanism and effectiveness of burying behaviours in the ocellate river stingray (*Potamotrygon motoro*). We hypothesized that to bury, rather than digging, movement of the body and fins in *P. motoro* generate flow beneath the pectoral disc to fluidize and suspend sediment onto the dorsal and exposed surface. Furthermore, we predicted that an increase in aspects of the kinematics of the body and fins would increase the sediment flows, to move more sediment and ultimately increase the extent of burying. Videography was used to explore the kinematics and evaluate the effectiveness by which *P. motoro* buries, and particle image velocimetry was used to describe the dynamics of the fluidized sediment, and accordingly, reveal the mechanism of burying. This study advances our understanding of the biology of stingrays in relation to life in a benthic environment, describing a novel approach to burying in a flattened fish with flexible pectoral fins and a rigid body axis, and it is relevant to underwater robotic designs for crypsis and station holding in a benthic environment, inspired by biomimicry.

## MATERIALS AND METHODS

### Animals and housing

All procedures were approved by the University of Calgary animal care committee, following Canadian Council on Animal Care guidelines. Four ocellate river stingrays (*Potamotrygon motoro*) (pectoral disc width 14.9 ± 0.410 cm S.E.M.; mass 155 ± 10.2 g S.E.M.) were purchased from a licenced supplier and transported to facilities at the University of Calgary. *P. motoro* were housed in a cylindrical holding tank (180 cm diameter × 70 cm height, approximately 1400 L) with flow-through freshwater at 27° C, pH of 6.5, bubbled with air. Room lighting was provided with a 12:12h on:off cycle. The stingrays were provided 5 cm^3^ of frozen bloodworms per ray once a day, and were monitored to ensure feeding. Furthermore, the stingrays were acclimated for two months prior to experiments, water chemistry was measured daily, and the fish had regular veterinary oversight.

### Data collection

A separate rectangular tank with dimensions 75 cm length × 32 cm width × 46 cm height was used for experimentation; water chemistry and temperature were the same as the holding tank. Aquarium substrate, with a maximum grain diameter of 1 mm (Crystal River Sediment, CaribSea, Fort Pierce, Florida, USA), was evenly spread across the bottom of the tank, at a depth of 7 cm. An individual ray was transferred to the experimental tank via a rubber mesh net. Burying behaviours were captured using three cameras filming at 120 fps at 720p (Hero3 Black, GoPro Inc, San Mateo, California, USA), one filming from the dorsal view and two filming across the substrate from each end of the tank. A measuring stick was placed on the bottom of the tank to calibrate the camera from the dorsal view, whereas measuring sticks were placed vertically at four different locations along the length of the tank and a linear regression relating distance from the camera to the filming location was used to calibrate cameras filming along the substrate of the tank. A pair of light emitting diodes attached to a plastic pipe submerged in the tank was used to synchronize the cameras immediately after the burying event was performed. A total of 20 responses were collected; each fish performed 5 burying events. The four stingrays were rotated into the experimental tank throughout the study, such that only a single burying event was recorded from an animal before it was placed back into the holding tank and another animal moved into the experimental tank.

### Data analysis

#### Sediment flow patterns and speed

Particle image velocimetry via MATLAB PIVlab 2.01 (Garcia, 2011; Thielicke, 2014; Thielicke and Stamhuis, 2014) was used to describe the dynamics of sediment movement during burying. From the anterior view, 10 burying events, where the head of the ray was oriented toward the camera, were analysed to explore the pattern of the displacement of the sediment granules from underneath the disc onto the dorsal surface. As the sediment moved across the dorsal surface, maximum speed and direction of the sediment flow was determined frame-by-frame in two-dimensions from the dorsal view for all 20 burying events, starting when the sediment moved from the pectoral fins towards the midline, and ending when the sediment reached the midline or when the ray came to a rest (i.e. if the sediment did not reach the midline). The 10 most rapid measures of sediment speed for each burying event were averaged and this average was used as sediment speed for statistical analysis. For both dorsal and anterior view analysis, the exposed body of the ray was masked for each frame such that PIV was measuring only sediment flow and not movement of the body (Fig. 1 & 2). Furthermore, any air bubbles present in the video frames were masked so they did not interfere with PIV. Three passes were conducted on the videos: one with 64×64 pixel windows, one with 32×32, and one with 16×16. Increasing the interrogation area with a fourth pass of 128×128 pixel windows did not yield different results in 5 videos analysed, and decreasing the interrogation area with a fourth pass of 8×8 pixel windows was too small relative to the size and displacement of the sediment particle, yielding substantial noise in the data.

**Fig. 1.**
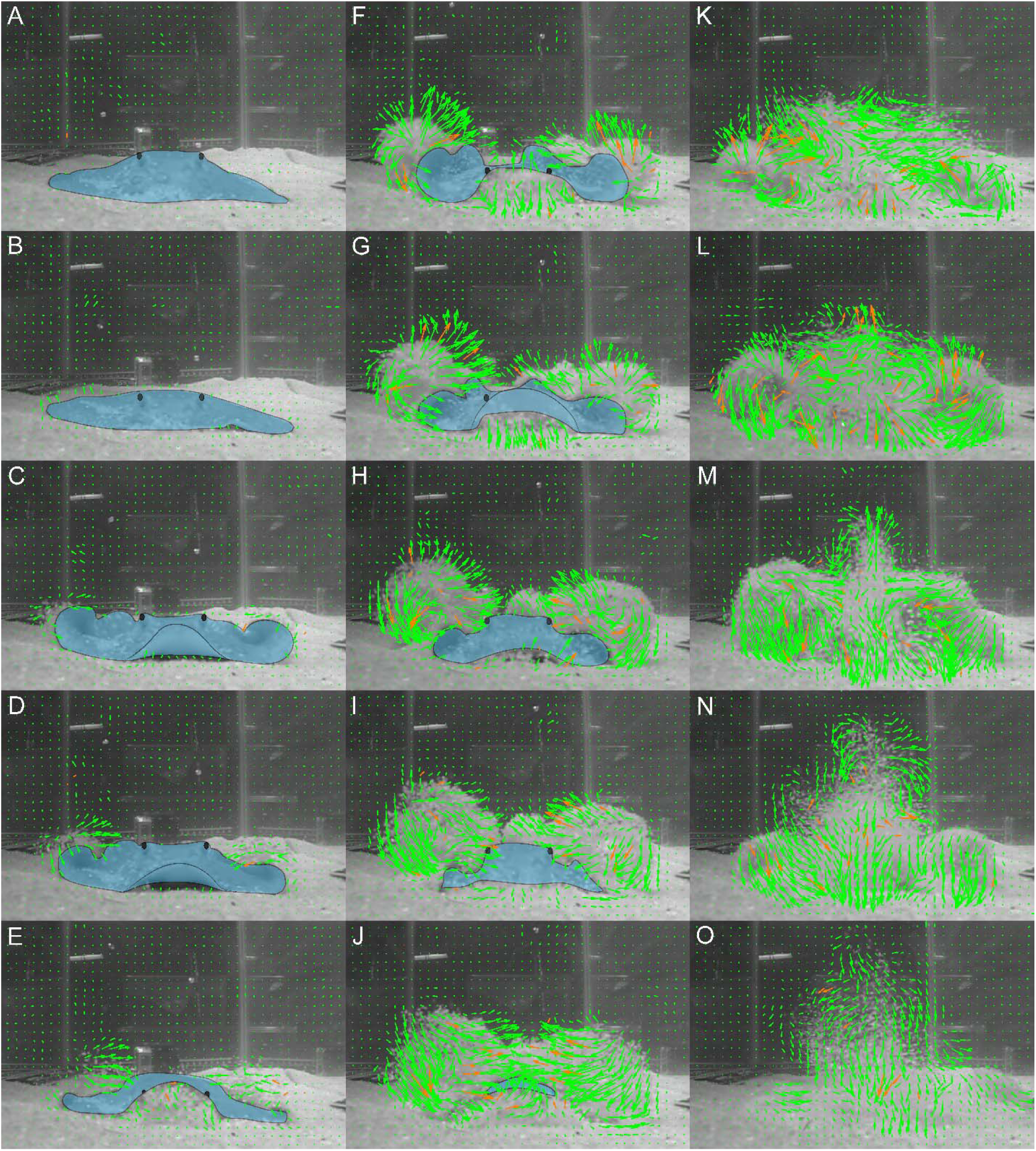
Particle image velocimetry analysis of a burying event, viewed from the anterior, showing the direction and relative speed of sediment granules. Granules are lifted as vortices, move along the ventral surface of the fins and are then shed towards the dorsal midline, where they collide and annihilate as indicated by the upwards jets. (A-B) From a resting position, the fish pressed its body downwards. (C-D) The rostrum lifted upwards, followed by the body pulling upwards, as the fins folded up and over. (E) The rostrum subsequently pressed onto the substrate as the body pushed downwards and the fins recovered towards the original position by rolling outwards. (F-G) Again, the rostrum and then the body pulled upwards, as the fins folded up and over suspending and directing vortices of fluidized sediment along the ventral surface of the fins towards the dorsal midline. (H-I) The vortices of sediment moved towards the midline, as the body presses downwards. (K-M) The vortices collided along the midline, as another set of vortices were formed by subsequent finbeat and directed towards the midline. (N) The second set of vortices collided over the midline. (O) The sediment dissipated and settled onto the fish. In all images, every second vector is shown for clarity. Orange vectors indicate data interpolated from neighboring vectors.

**Fig. 2.**
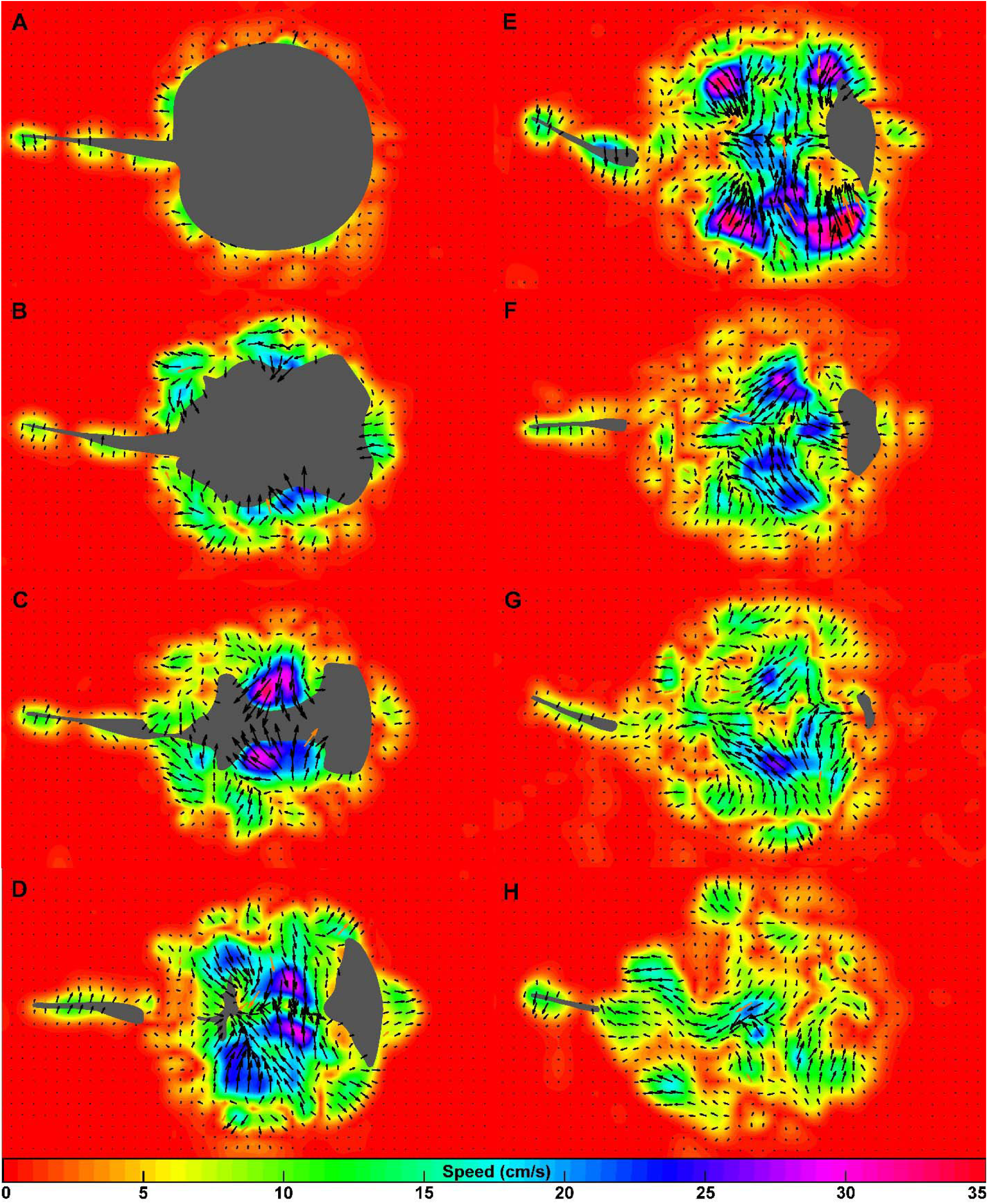
Particle image velocimetry analysis of a burying event, viewed from the dorsal surface, showing the measured speed and direction of sediment moving across the dorsal surface of the fish towards the sagittal midline, where the sediment from each fin collides and generates anteroposterior jets of sediment flow that reorient the sediment towards the head and tail. (A-C) The fins folded up and over, suspending and directing sediment towards the midline. (D-F) The sediment moved towards the midline and collided, changing direction towards the head and tail base, as another surge of sediment was suspended and directed towards the midline for another collision by a subsequent finbeat. (G-H) The sediment settled over the dorsal surface of the fish. In all images, every second vector is shown for clarity. Orange vectors indicate data interpolated from neighboring vectors. Color bar scale at the bottom of the image indicates particle speed.

#### Kinematics

Digital video images were analysed using ImageJ. Using images from the camera filming an anterior view of the ray along the substrate, the location of an eye was tracked for successive frames to investigate the kinematics of the down-up pumping motions (i.e. oscillations) exerted by the body along the vertical axis relative to the substrate. The number of body pumps was defined as the number of down-up oscillations during the burying event. The oscillations of the body were not symmetrical, and therefore, the downward motions were defined as a push (i.e. toward the substrate), whereas the upward motions were defined as a pull (i.e. away from the substrate), and displacement and speed for pushing and pulling motions were calculated separately. Displacement was defined as the cumulative change in the vertical position of the eye for a given push or pull, and the maximum and average displacement of the body pumping motions throughout burying were measured. Speed was calculated using the two-dimensional change in the vertical position of the eye over time between successive frames, and the average speed of the body pumping motions were measured in addition to the maximum speed. The onset of the body pumping motions (always a push) was defined as the point at which push speed was 10% of the maximum of the initial push speed. Average body-pump frequency was defined as the total number of oscillations divided by the duration (in seconds) of these oscillations. Body pumping often began before motion of the pectoral fins, and therefore, the body pump oscillations that were performed during the finbeats were selected to evaluate the relationship between body pump frequency and finbeat frequency, defined below.

As *P. motoro* buried, the pectoral fins rapidly and repeatedly folded up and over, and the three-dimensional bending during burying events appeared far more complex than what has been described for routine swimming (Blevins and Lauder, 2012). We anticipated that the lateral motion of the fins towards the sagittal midline would most impact the two-dimensional speed of sediment as it moved across the dorsal surface. Hence, analysis of fin motions was simplified to only two dimensional movements as measured from the dorsal view, whereby the length of a line transecting the centre of the disc from the lateral edge of the dextral fin to the lateral edge of the sinistral fin was measured for successive frames and used as a proxy for the displacement and speed of fin movement. Finbeats were assessed until the fins either came to rest or the lateral edges of the fins were covered by substrate. The number of finbeats was defined as the number of oscillations of the fins folding over towards the sagittal midline, and then recovering to the original position, during a burying event. The average finbeat frequency was defined as the number of oscillations divided by the duration of these oscillations. Finbeat displacement was defined as the cumulative two-dimensional change in length of the line from fin tip-to-tip, and the maximum and average displacements of the finbeats throughout the burying event were measured. Finbeat speed was calculated using the two-dimensional change in length of the line from tip-to-tip between successive frames, and the maximum and average speed of the finbeat motions towards the midline were measured. Displacement and speed values were divided by two, to provide values that were associated with one fin, and the onset of finbeat motions was defined as the point where finbeat speed was 10% of the maximum for the initial finbeat speed.

#### Performance

Burying duration was defined as the onset of the body pumping motions to when the body pumping came to rest or the eyes were covered by substrate. Burying depth was measured as the displacement of the eye from its resting position immediately before burying commenced, to its position after the burying event occurred or when the eye was covered by substrate. Sediment coverage of the ray, from the dorsal view, was measured to compare the extent of burying of the head, body, fins and tail, and the effects of body kinematics on extent of coverage. From the dorsal view, a grid was created using ImageJ over the animal at rest before burying motions began, to define the surface area covered by sediment of different locations on the disc (Fig. 3). The outline of the ray was traced, and then points were marked around the lateral edge, separated by 30 degrees relative to the centre of the disc, starting from the rostrum, which was defined as 0 degrees. Then, anteroposterior and lateral transects were created to connect the points directly across from one another, creating a grid over the dorsal surface. Furthermore, the tail was divided into proximal (i.e. tail base proximal to the stinger) and distal (i.e. tail tip from the stinger outward) locations. This grid defined 18 different locations on the dorsal surface of the pectoral disc and tail (Fig. 3). The grid was then placed over the image of the animal once the burying event was completed, using the location of the eyes and the tail as a reference, which remained recognizable even when the rest of the ray was fully buried. For each location in the grid, the surface area covered by sediment was measured after burying. The percent of sediment coverage for each location was defined as the surface area of the location covered divided by the total surface area of that location, multiplied by 100. Additionally, the total sediment coverage of the ray (i.e. pectoral disc and tail) was defined as the sum of the amount of surface area covered for each location divided by the total surface area of the pectoral disc, multiplied by 100.

**Fig. 3.**
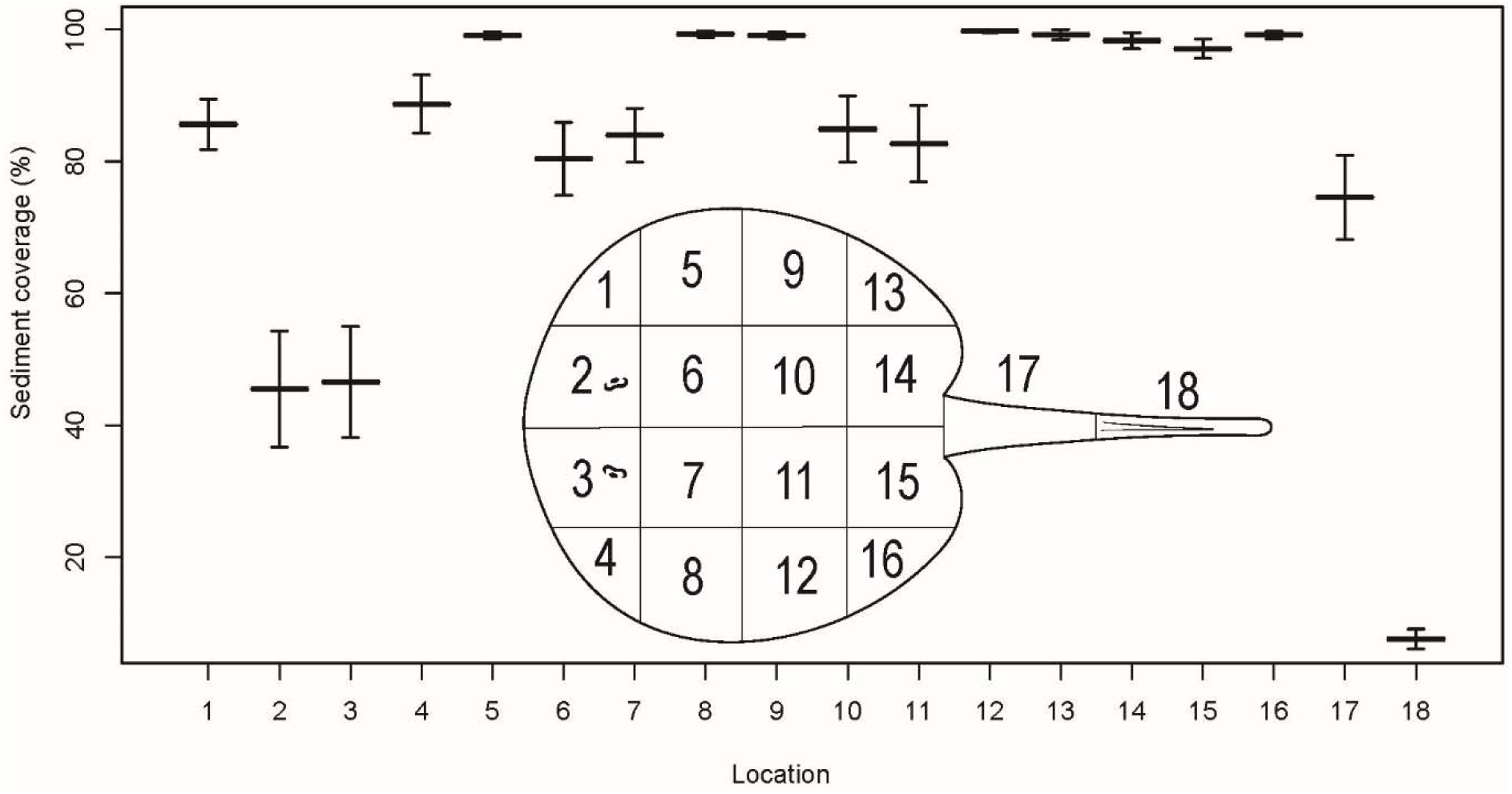
Sediment coverage of the different locations on the dorsal surface of the stingray. Mean and standard error of the mean. See table 1 for statistical comparisons.

#### Statistical analysis

All statistical analyses were performed using R software and accounted for repeated measures (R Development Core Team, 2008). A linear regression was used to test the relationship of sediment speed and sediment coverage. Data of sediment coverage was transformed using the arcsine of the square root of the percentage prior to statistical analysis. Linear regressions were also used to test the relationship between the frequency of the body pump and finbeat motions, and the impact of the following measures on sediment speed: number of finbeats, number of body pumps, average finbeat frequency, average body-pump frequency, maximum and average displacement and speed of the finbeats and the push and pull of the body pumping. Furthermore, linear regressions were used to test the relationship of burying depth and duration on sediment coverage. Moreover, a one-way ANOVA and Tukey HSD Post-Hoc Test was used to test for differences in sediment coverage of the different individuals, in addition to the different locations of the pectoral disc.

**Table 1.**
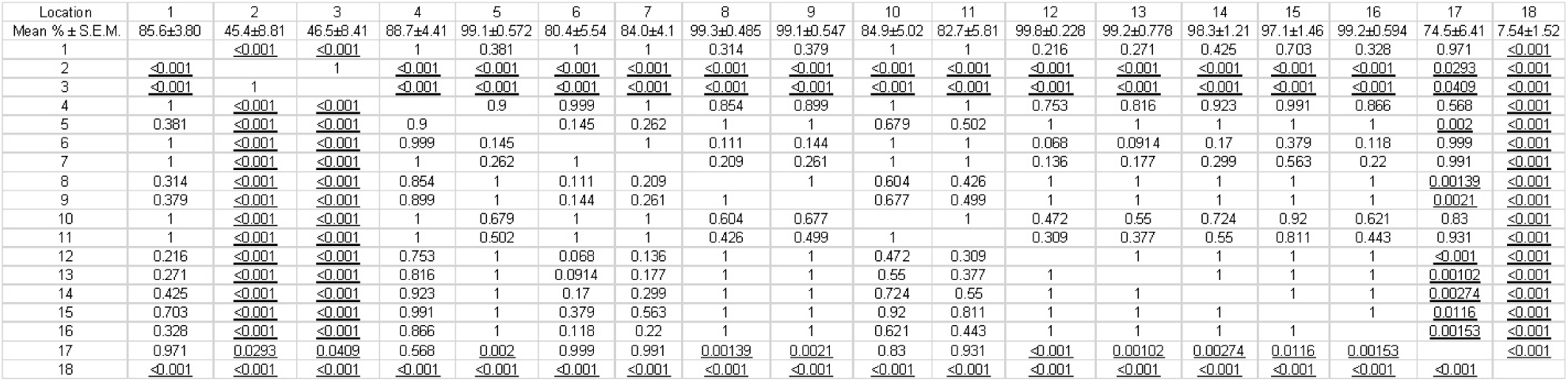
P-value statistics for comparisons in sediment coverage of the different locations on the dorsal surface of *P. motoro* (see Fig. 3 for locations). Row 1 represents the average sediment coverage for each location.

## RESULTS

### Sediment flow patterns

To bury, *P. motoro* repeatedly pumped the body up and down as the pectoral fins folded up and over (Fig. 1). The body pumping appeared to draw water beneath the rostrum and into the space between the body and substrate as the anterior portion of the pectoral fins pressed into the substrate, indicated by movement of sediment being sucked towards the rostrum (Fig. 2). The flow and pumping motions suspended sediment beneath the pectoral disc, and the movement of the fins generated and directed vortices of suspended sediment along the ventral surface of the fins as they were lifted. The vortices moved toward the fin tips, and then over and onto the dorsal surface of the body as the fins were depressed again, shedding the vortices of sediment over the dorsal surface of the body toward the midline where the sediment then settled. These patterns of sediment flow were resolved using video images and particle velocimetry analysis from both anterior and dorsal views.

From the anterior view, *P. motoro* lifted the sediment as vortices along the ventral surface of the pectoral fins, such that a circular, rotating mass of sediment particles translated in the same direction as the movement of fins, while the particles rotated in a direction counter to the dorso-medial movement of the tips of the fins (Fig. 1). The vortices then rolled over the tips of the fins as the fins were depressed, and were shed across the dorsal surface where they dissipated. The fins subsequently recovered into the original position by rolling the tips away from the body midline, laterally and downwards along the dorsal surface of the fish, so that the fin tips were maintained in close proximity to the dorsal surface and thus moved underneath the suspended sediment during the recovery stroke. In 5 of the trials, from the anterior view, the sediment vortices that were shed from the two fins collided over the dorsal midline of the fish, where they annihilated and upward vertical jets of sediment were produced as a result, before the sediment settled onto the dorsal surface (Fig. 1). From the dorsal view, the sediment was shed primarily along the region of the pectoral fins approximately posterior to the eye and anterior to the tail, in a lateral direction towards the longitudinal midline (Fig. 2). The velocity profiles suggest the sediment settled first onto the fins, and then onto the body (Fig. 3). In the 5 trials where the sediment shed from each fin collided along the midline, immediately following collision jets of high sediment velocity were created in the anteroposterior direction, resulting in sediment moving toward the head and the tail base, well anterior and posterior to the initial vortices of sediment that were shed over the dorsal surface by the fins (Fig. 2).

### Kinematics, sediment speed and performance

The mean sediment coverage of all burying events was 82.5% ± 3.0 S.E.M, and ranged from 60.4 to 98.1 % (Fig. 4). The posterior portion of the pectoral fins, which were those involved in the finbeats (location 5, 8, 9, 12, 13 and 16), were covered more than the fins anterior to the eye (location 1 and 4), the head (location 2 and 3), the body (location 6, 7, 10 and 11) and the tail base and stinger (location 17 and 18) (Fig. 3 & Table 1). The fins anterior to the eye, the body, and the tail base were covered more than the head and the stinger of the tail. The head was covered more than the stinger of the tail. Sediment coverage was significantly greater in ray 1 and 2 when compared to ray 4; otherwise, mean sediment coverage was not statistically different between individuals (1 vs 2, p = 0.216; 1 vs 3, p = 0.1; 1 vs 4, p < 0.001; 2 vs 3, p = 0.969; 2 vs 4, p = 0.0284; 3 vs 4, p = 0.0678) (Fig. 4).

**Fig 4.**
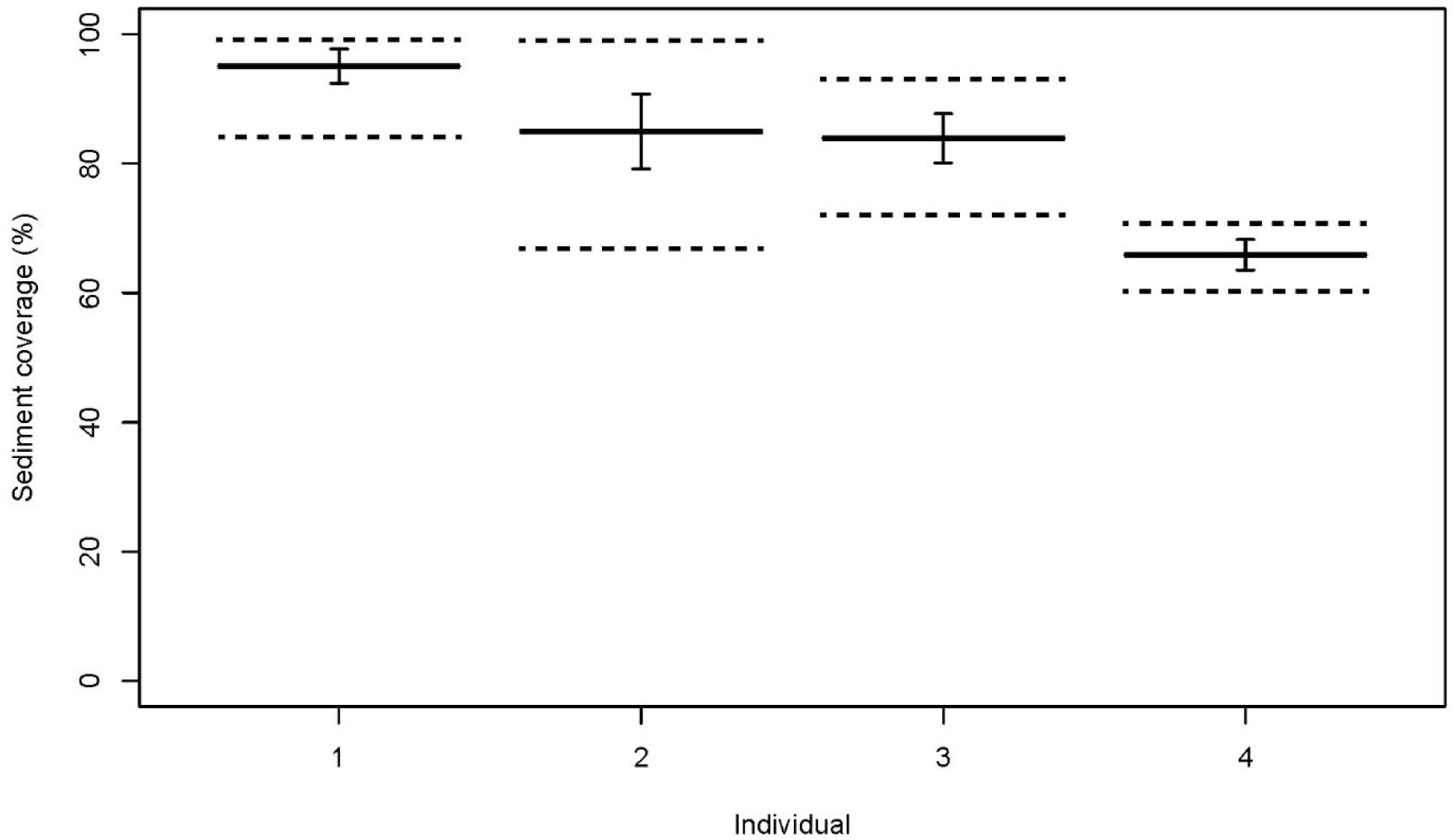
Average total sediment coverage of the four individuals. Vertical bars indicate the standard error of the mean, and horizontal dashed lines represent the ranges of coverage observed across the 5 burying events for each animal (ray 1: 95.1% ± 2.23 S.E.M.; ray 2: 85.0% ± 2.24 S.E.M.; ray 3: 83.9% ± 3.81 S.E.M.; ray 4: 65.9 ± S.E.M.).

Body pumping motions were often initiated before finbeat motions (65% of trials). As the frequency of the body pumping increased, frequency of the finbeats also increased (p < 0.001; Fig. 5), and the slope of the relationship between these parameters was not different than 1, indicating these motions occurred in synchrony and suggesting coordination of sediment fluidization beneath the body and the movement of that sediment by the fins onto the dorsal surface. The body pump and finbeat motions were coordinated such that maximum finbeat speed and maximum body-pull speed were closely aligned, as determined from the measured time course of these movements, whereas maximum body-push speed occurred during the recovery stroke of the fins. It was anticipated that the vigor of the body and fin motions should relate to the extent of sediment movement, and so relationships between displacement, speed and frequency of body and fin movements and sediment movement were explored. The number of body pumps (p = 0.644) and finbeats (p = 0.677), and the average body pump frequency (p = 0.072) and finbeat frequency (p = 0.0763), did not significantly impact the speed of the sediment as it moved towards the midline of the fish (Fig. 5). Conversely, as the maximum and average displacement of the body push (p < 0.001 & p < 0.001, respectively), body pull (p < 0.001 & p < 0.001, respectively), and maximum and average speed of body push (p < 0.001 & p < 0.001, respectively) and body pull (p < 0.001 & p < 0.001, respectively) increased, sediment speed increased (Fig. 6). Repeated measures of individuals influenced the positive relationship between the average pull displacement and sediment speed; ray 2 and 3 individually showed a positive relationship between the two parameters, whereas ray 1 and 4 individually showed no significance. Moreover, as the maximum and average speed of finbeats (p < 0.001 & p < 0.001, respectively) and finbeat displacement (p < 0.001 & p = 0.00202, respectively) increased, sediment speed increased (Fig. 7). Furthermore, as sediment speed increased, sediment coverage of the dorsal surface increased (p < 0.001; Fig. 8). Moreover, burying duration did not have an impact on sediment coverage (p = 0.572), and as burying depth increased the extent of sediment coverage increased (p < 0.001) (Fig. 8).

**Fig. 5.**
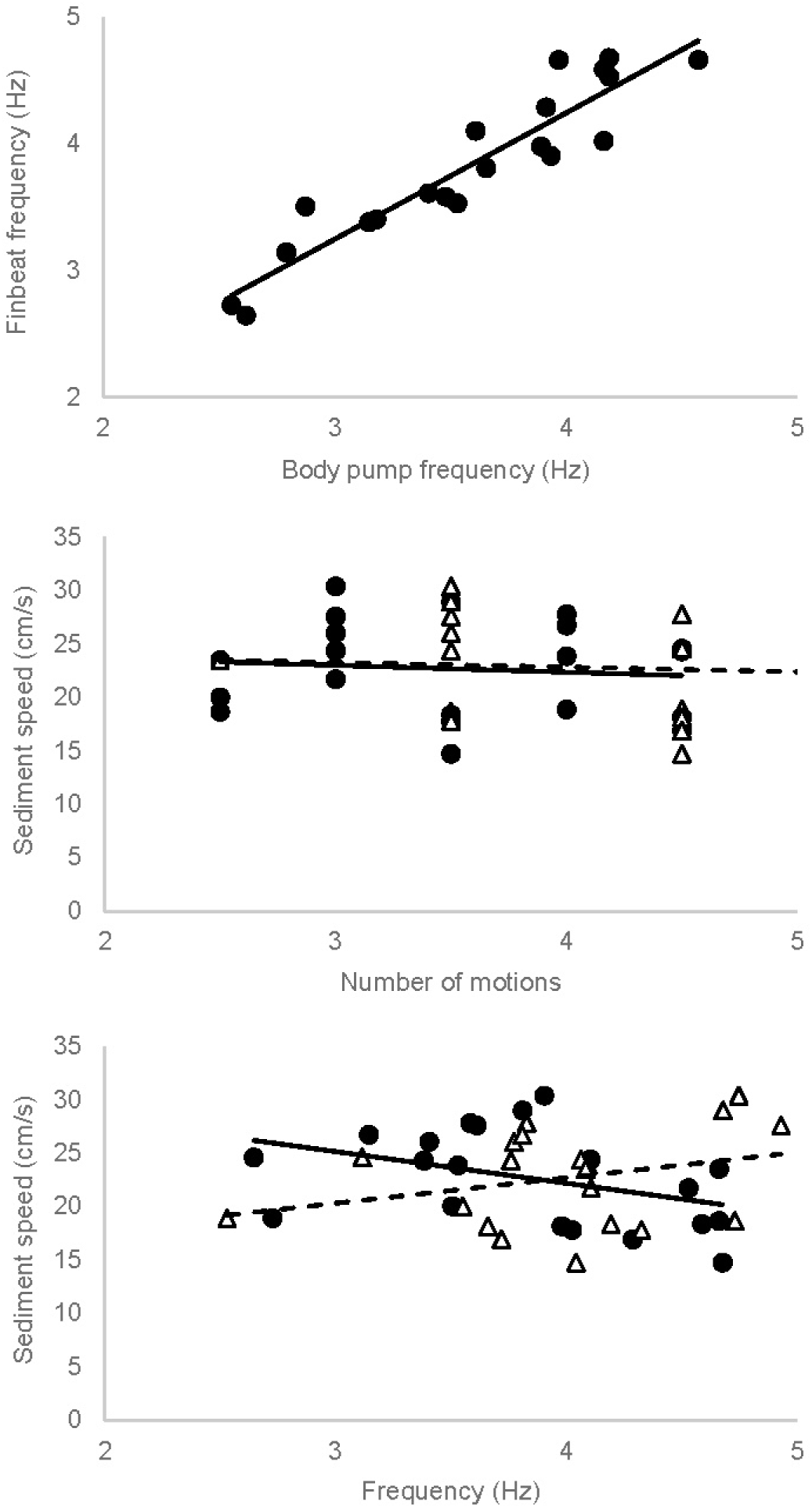
A) The relationship between body pump frequency and finbeat frequency (p < 0.001, R^2^ = 0.86, y = 1.0 × + 0.26). B) The relationship between the number of finbeats or body pumps and sediment speed during a burying event. The black circles and solid line represent finbeat number (p = 0.67, R^2^ = −0.045, y = −0.65x + 25) and the white triangles and dashed line represent body pump number (p = 0.64, R^2^ = −0.043, y = −0.46x + 25). C) The relationship between average finbeat or body pump frequency and sediment speed. The black circles and solid line represent finbeat frequency (p = 0.076, R^2^ = 0.12, y = −3.0x + 34) and the white triangles and dashed line represent body pump frequency (p = 0.072, R^2^ = 0.13, y = 2.4x + 13).

**Fig. 6.**
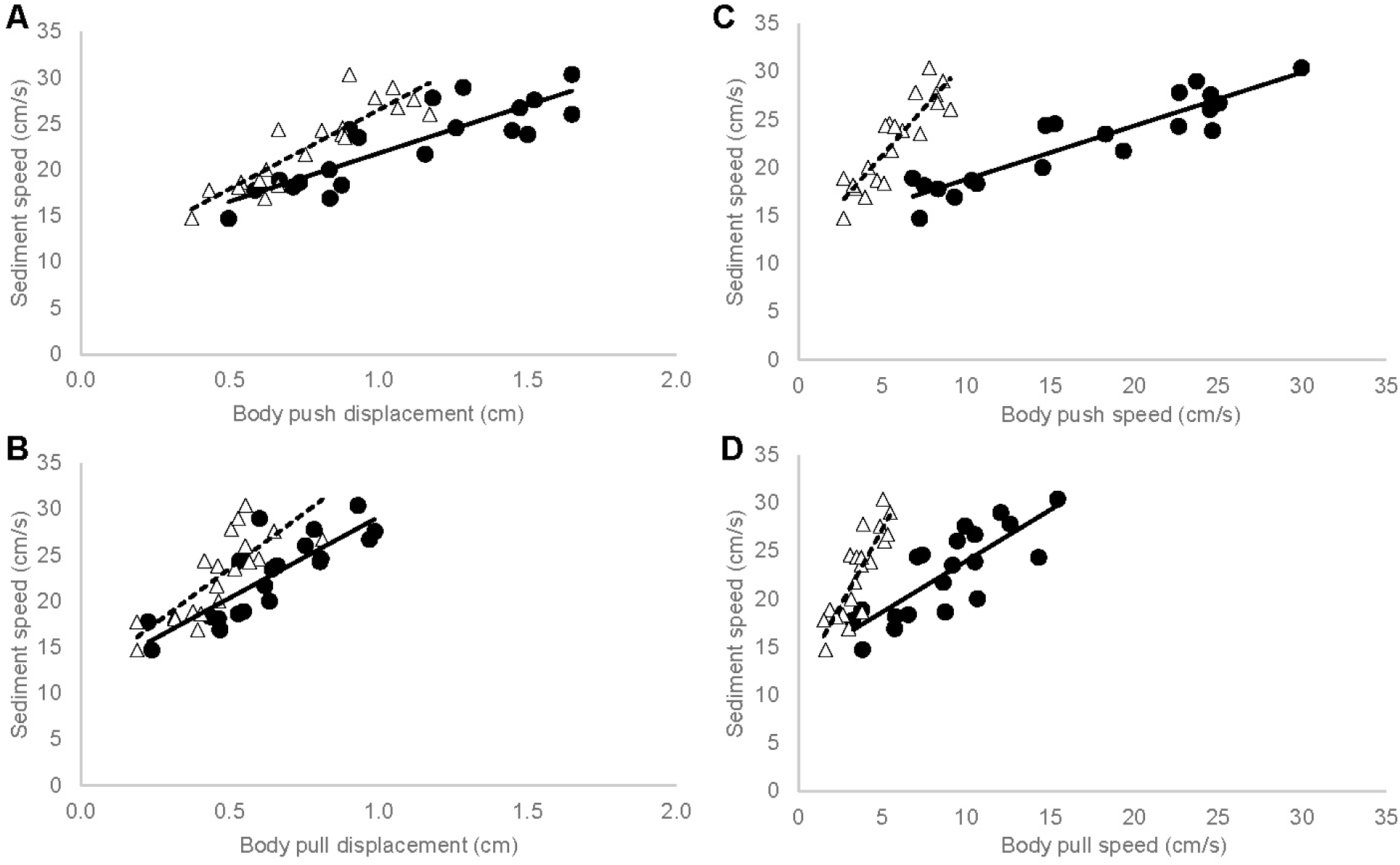
A) The relationship between body-push displacement and sediment speed. The black circles and solid line represent maximum push displacement (p < 0.001, R^2^ = 0.73, y = 10x + 11), and the white triangles and dashed line represent average push displacement (p < 0.001, R^2^ = 0.77, y = 17x + 9.3). B) The relationship between body-pull displacement and sediment speed. The black circles and solid line represent maximum pull displacement (p < 0.001, R^2^ = 0.68, y = 18x + 12), and the white triangles and dashed line represent average pull displacement (p < 0.001, R^2^ = 0.61, y = 24x + 12). C) The relationship between body-push speed and sediment speed. The black circles and solid line represent maximum push speed (p < 0.001, R^2^ = 0.83, y = 0.56x + 13), and the white triangles and dashed line represent average push speed (p < 0.001, R^2^ = 0.77, y = 2.0x + 11). D) The relationship between body-pull speed and sediment speed. The black circles and solid line represent maximum pull speed (p < 0.001, R^2^ = 0.62, y = 1.1x + 13), and the white triangles and dashed line represent average push speed (p < 0.001, R^2^ = 0.77, y = 3.2x + 11).

**Fig. 7.**
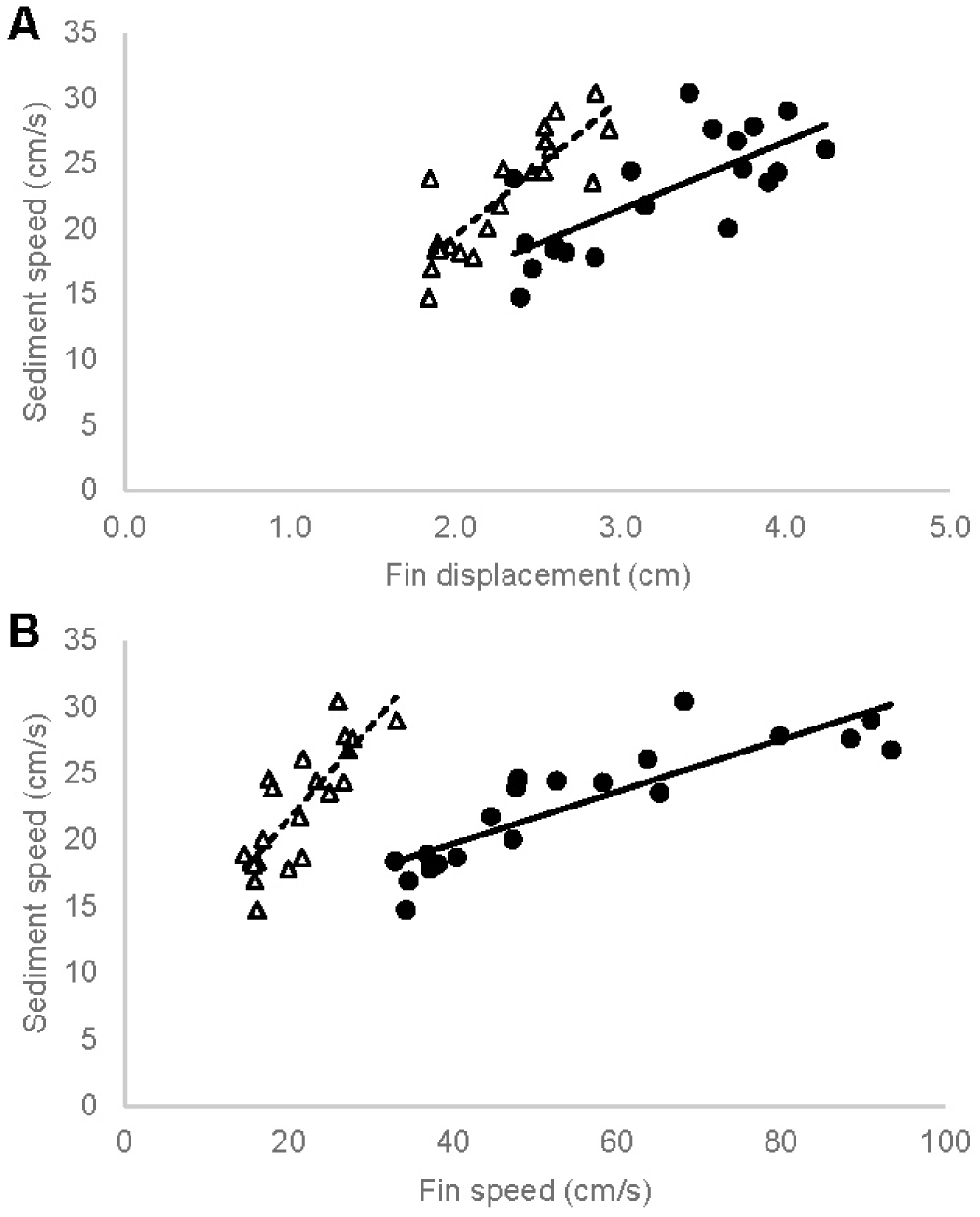
A) The relationship between fin displacement and sediment speed. The black circles and solid line represent maximum fin displacement (p < 0.001, R^2^ = 0.52, y = 5.2x + 5.8) and the white triangles and the dashed lines represent the average fin displacement p <0.001, R^2^ = 0.68, y = 10x −1.3). B) The relationship between fin speed and sediment speed. The black circles and solid line represent maximum fin speed (p<0.001, R^2^ = 0.74, y = 0.20x + 12), and the white triangles and dashed lines represent average fin speed (p < 0.001, R^2^ = 0.64, y = 0.70x + 7.6).

**Fig. 8.**
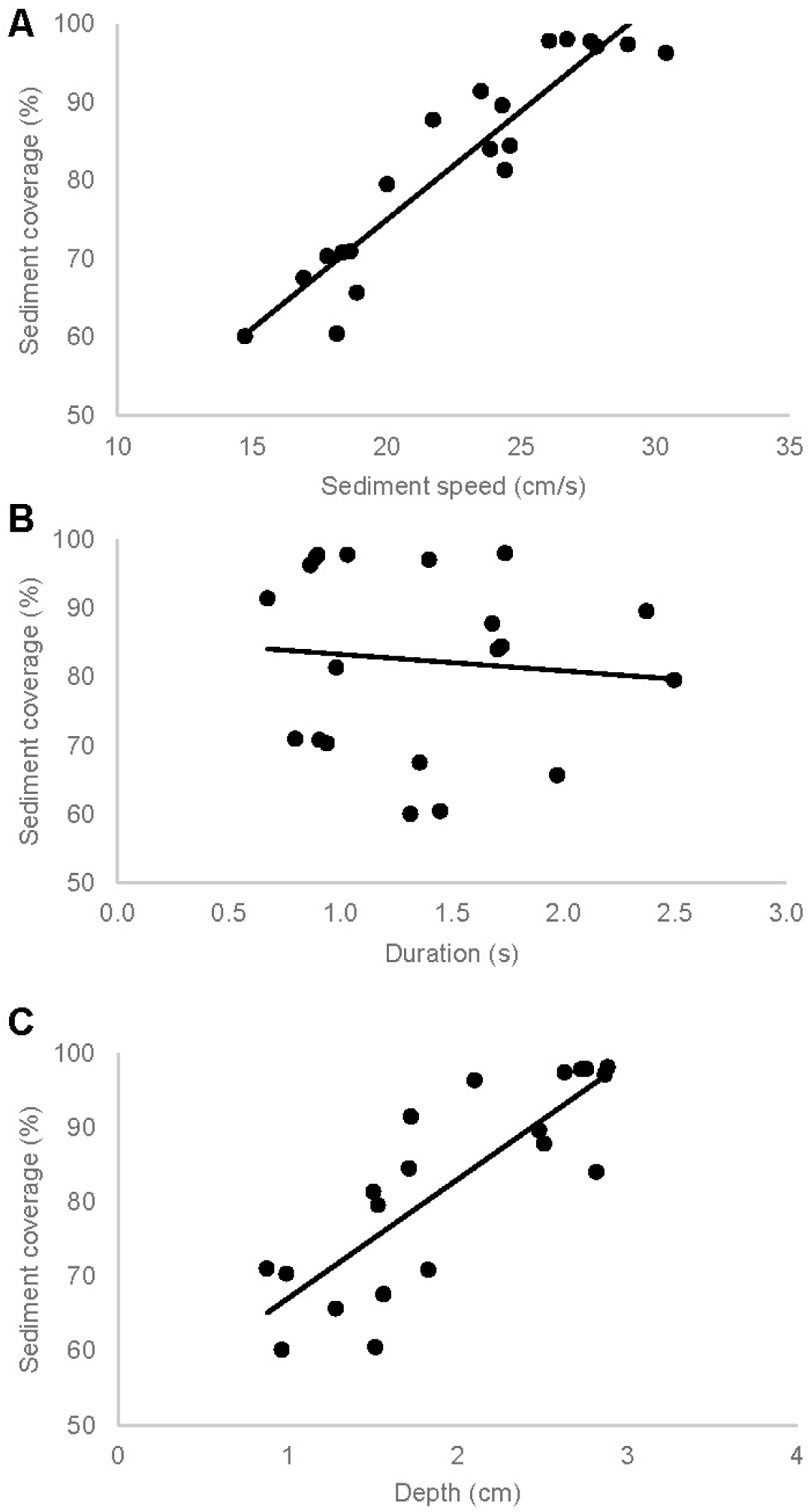
A) The relationship between sediment speed and sediment coverage (p < 0.001, R^2^ = 0.83, y = 2.8x + 19). B) The relationship between sediment coverage and duration of burying (p = 0.57, R^2^ = −0.037, y = −2.4x + 85). C) The relationship between sediment coverage and burying depth (p < 0.001, R^2^ 0.68, y = 16x + 51).

## DISCUSSION

This study revealed that the mechanism of burying employed by *P. motoro* permits effective control of sediment vortices and flows to modulate the extent of burying. Rather than digging, the downwards-pushing and upwards-pulling movement of the body of *P. motoro*, coupled with movement of the pectoral fins including around the rostrum to form and release a seal along the substrate, functioned like a piston with valves to control the flow of water and generate changes in pressure that fluidized and suspended sediment beneath the ray. As the fins folded up and over, vortices of sediment were elevated upwards and along the ventral surface of the fins, transferred across the lateral edge of the fins toward the dorsal surface, and then were shed medially where they dissipated over the dorsal side of the fish, as the fins then moved under the vortices of sediment and recovered to the original position (Fig. 1). In the most vigorous burying events, the vortices of sediment travelled toward the dorsal midline where they collided and were annihilated, resulting in jets of sediment being directed upwards along the vertical axis, and forwards and backwards along the anteroposterior axis of the animal (Figs. 1 & 2). This led to near complete coverage of the ray with sediment, including the head and most of the tail except for the stinger, despite only using about the posterior two-thirds of the pectoral fins to lift sediment from the substrate. Following is a description of how the body and fin movements appear to generate these patterns of sediment dynamics and impact the extent of burying.

The movements of *P. motoro* and the sediment suggest that stingrays rely on changes in pressure underneath the pectoral disc, particularly suction, to fluidize sediment beneath the disc and move it along the ventral surface of the fins. Water from the surrounding environment was drawn into the cavity beneath the animal through suction pressure created as the body pulled upwards, flowing in via a tunnel created by the ray raising the rostrum off the substrate while the anterior portion of the pectoral fins maintained contact with the substrate, similar to the mechanisms employed by stingrays to feed (Wilga et al., 2012). In support of this mechanism, granules of sediment near the rostrum were often observed to be sucked in under the disc as the rostrum and body lifted upwards (Fig. 2). Moreover, an increase in pressure beneath the pectoral disc most likely occurred as the body subsequently pushed towards the substrate and the rostrum and pectoral fins were pressed against the substrate, expelling the fluidized sediment toward the lateral edges of the pectoral disc. Furthermore, the upward motion of the fins generated suction along the ventral side of the fins, drawing the fluidized sediment from underneath the ray and transferring it along the ventral and presumably low pressure surface of the fins as vortices, similar to the appearance of hydrodynamic vortices formed and shed along the trailing surface of the caudal fin during stage 1 of the C-start in fishes (Borazjani et al., 2012; Tytell and Lauder, 2008) (Fig. 1).

Variation in the relationships between several aspects of the kinematics, the speed of sediment movement, and sediment coverage suggests that the mechanism employed by *P. motoro* to bury can control the speed of sediment flow and thus the extent of burying. When *P. motoro* increased the speed and displacement of the body pumping and finbeat motions, this increased the speed of the sediment movement (Fig. 6 & 7) and fluidized more sediment to increase sediment coverage of the dorsal surface (Fig. 8). An increase in speed and displacement as the body pulled upwards away from the substrate most likely generated a greater magnitude of suction beneath the disc, and hence, a greater rate of water flow in through the tunnel beneath the rostrum, similar to what is observed when stingrays increase suction in feeding (Wilga et al., 2012). This would likely fluidize more sediment beneath the body. Furthermore, an increase in the displacement and the speed of the body subsequently pushing downwards would likely increase the pressure beneath the disc and expel more of the fluidized sediment toward the periphery, and perhaps this increased flow might also loosen and fluidize more sediment as well. The capacity to expel more sediment would also allow the fish to burry deeper in the substrate (Fig. 8). The frequency and number of body and fin movements was not related to burying (Fig. 5), nor was the duration of the burying event (Fig. 8), and the average frequency of the body pumping and finbeat motions remained within a narrow range in all individuals as they buried. This is opposed to observations in burying flatfish, that increase sediment coverage by increasing the frequency of body undulations (Corn et al., 2018; McKee et al., 2016). In *P. motoro*, increasing the number of body and finbeat movements might move more sediment in total, but would not necessarily increase the speed and thus distance that the sediment travels, and therefore, would have less impact on extent of burying. And changes in the frequency of movements was more a reflection of changes in the duration of pauses between movements rather than the rate of movement itself, which again would have relatively little impact on the speed and distance that sediment would move. Therefore, it appears burying in *P. motoro* is not necessarily related to how long the ray partakes in the behaviour (duration of burying even, number of fin beats), rather it is dependent on the vigour of the burying event. Moreover, it is not clear what motivated the variation in sediment coverage of *P. motoro* within individuals. Individuals demonstrated different levels of sediment coverage when compared to one another, whereby one of the rays tended to only bury its pectoral fins leaving most of the body and the head unburied for all burying events, and another ray never buried its head fully (Fig. 4). Hence, this suggests different thresholds or motivations to bury, or perhaps limitation in physical capability of individuals might exist in stingrays.

There was also evidence that stingrays can selectively control sediment flows and ultimately what specific parts of the dorsal surface were covered. *P. motoro* did not always cover the entire dorsal surface with sediment (Fig. 3), and the pattern of how the sediment moved up and over onto the dorsal side of the pectoral disc impacted the sediment coverage of different locations on the fish. Sediment initially moved onto the pectoral disc mostly along the portion of the fins that were involved in finbeats, which was predominantly the fins posterior to the eye (Fig. 2). The fins anterior to the eye tended to remain close to the substrate, contributing to forming the tunnel through which water was drawn under the body as the rostrum lifted upwards, but not contributing greatly to the finbeat motions that directed sediment onto the dorsal surface (Fig. 1 & 2). The rostrum pressed down onto the substrate, but it did not tend to flick sediment onto the surface as it lifted away from the substrate. As such, little substrate moved onto the pectoral disc across the rostrum and anterior portion of the pectoral fins. Likewise, the tail region was not involved in moving sediment onto the dorsal surface. When the speed and displacement of the body and fin kinematics were relatively low, the vortices of sediment did not reach the midline before they dissipated, and consequently the sediment settled only on the pectoral fins, and only in the middle region of the fish. As the speed and displacement of the body pumping and finbeats became faster, sediment moved across the posterior fins and towards the sagittal midline, and settled on the fins and much of the body, but again primarily the posterior portion. When the speed and displacement of the finbeats were greatest, the vortices of sediment from opposite sides of the body converged above the body midline, and the pathway of the sediment flow then shifted from a lateral orientation to an anteroposterior and vertical orientation, producing jets of sediment towards the body and the tail and upwards (Fig. 1 & 2). The anteroposterior and upwards jets of sediment suggest that stingrays might be exploiting the collision and annihilation of vortices to cover the head and the tail base, where energy in the colliding and collapsing vortices is redirected radially and axially (Kudela and Kosior, 2014; Lim and Nickels, 1992).

## ACKNOWLEDGEMENTS

This research was supported by an NSERC Discovery Grant to DAS, and University of Calgary Killam and NSERC Alexander Graham Bell Scholarships to SGS. We would like to thank Josh D. Wilson for his help with coding in R.

## REFERENCES

Allen, L. G. and Pondella, D. J. (2006). Surf Zone, Coastal Pelagic Zone, and Harbors. In The Ecology of Marine Fishes California and Adjacent Waters (ed. Allen, L. G.), Pondella, D. J.), and Horn, M. H.), pp. 149–166. Berkeley: University of California Press.

Aschliman, N. C., Nishida, M., Miya, M., Inoue, J. G., Rosana, K. M. and Naylor, G. J. P. (2012). Body plan convergence in the evolution of skates and rays (Chondrichthyes: Batoidea). Mol. Phylogenet. Evol. 63, 28–42.

Blevins, E. L. and Lauder, G. V. (2012). Rajiform locomotion: three-dimensional kinematics of the pectoral fin surface during swimming in the freshwater stingray *Potamotrygon orbignyi*. J. Exp. Biol. 215, 3231–3241.

Borazjani, I., Sotiropoulos, F., Tytell, E. D. and Lauder, G. V. (2012). Hydrodynamics of the bluegill sunfish C-start escape response: three-dimensional simulations and comparison with experimental data. J. Exp. Biol. 215, 671–684.

Corn, K. A., Farina, S. C., Summers, A. P. and Gibb, A. C. (2018). Effects of organism and substrate size on burial mechanics of English sole, *Parophrys vetulus*. J. Exp. Biol. 221, jeb176131.

Fontanella, J. E., Fish, F. E., Barchi, E. I., Campbell-Malone, R., Nichols, R. H., DiNenno, N. K. and Beneski, J. T. (2013). Two- and three-dimensional geometries of batoids in relation to locomotor mode. J. Exp. Mar. Bio. Ecol. 446, 273–281.

Franklin, O., Palmer, C. and Dyke, G. (2014). Pectoral fin morphology of batoid fishes (Chondrichthyes: Batoidea): Explaining phylogenetic variation with geometric morphometrics. J. Morphol. 275, 1173–1186.

Garcia, D. (2011). A fast all-in-one method for automated post-processing of PIV data. Exp. Fluids 50, 1247–1259.

Gibson, R. N. and Robb, L. (1992). The relationship between body size, sediment grain size and the burying ability of juvenile plaice, *Pleuronectes platessa* L. J. Fish Biol. 40, 771–778.

Gidmark, N. J., Strother, J. A., Horton, J. M., Summers, A. P. and Brainerd, E. L. (2011). Locomotory transition from water to sand and its effects on undulatory kinematics in sand lances (Ammodytidae). J. Exp. Biol. 214, 657–664.

Goes de Araújo, M., Charvet-Almeida, P., Almeida, M. P. and Pereira, H. (2004). Freshwater stingrays (Potamotrygonidae): status, conservation and management challenges. Inf. Doc. AC 20, 1–6.

Kudela, H. and Kosior, A. (2014). Numerical study of the vortex tube reconnection using vortex particle method on many graphics cards. J. Phys. Conf. Ser. 530,.

Lim, T. T. and Nickels, T. B. (1992). Instability and reconnection in the head-on collision of two vortex rings. Nature 357, 225–227.

McKee, A., MacDonald, I., Farina, S. C. and Summers, A. P. (2016). Undulation frequency affects burial performance in living and model flatfishes. Zoology 119, 75–80.

Parson, J. M., Fish, F. E. and Nicastro, A. J. (2011). Turning performance of batoids: Limitations of a rigid body. J. Exp. Mar. Bio. Ecol. 402, 12–18.

Rosenberger, L. (2001). Pectoral fin locomotion in batoid fishes: undulation versus oscillation. J. Exp. Biol. 204, 379–394.

Rosenberger, L. and Westneat, M. (1999). Functional morphology of undulatory pectoral fin locomotion in the stingray *Taeniura lymma* (Chondrichthyes: dasyatidae). J. Exp. Biol. 202 Pt 24, 3523–39.

Schaefer, J. T. and Summers, A. P. (2005). Batoid wing skeletal structure: Novel morphologies, mechanical implications, and phylogenetic patterns. J. Morphol. 264, 298–313.

Seamone, S. G., McCaffrey, T. M. and Syme, D. A. (2019). Disc starts: the pectoral disc of stingrays promotes omnidirectional fast starts across the substrate. Can. J. Zool. 605, 597–605.

Stoner, A. W. and Ottmar, M. L. (2003). Relationships between size-specific sediment preferences and burial capabilities in juveniles of two Alaska flatfishes. J. Exp. Mar. Bio. Ecol. 282, 85–101.

Tanda, M. (1990). Studies on burying and ability in sand and selection to the grain size for hatchery-reared marbled sole and Japanese flounder. Nippon Suisan Gakkaishi 56, 1543–1548.

Tatom-Naecker, T. M. and Westneat, M. W. (2018). Burrowing fishes: kinematics, morphology and phylogeny of sand-diving wrasses (Labridae). J. Fish Biol. 860–873.

Thielicke, W. (2014). The flapping flight of birds - analysis and application.

Thielicke, W. and Stamhuis, E. J. (2014). PIVlab – towards user-friendly, affordable and accurate digital particle image velocimetry in MATLAB. J. Open Res. Softw. 2,.

Tytell, E. D. and Lauder, G. V. (2008). Hydrodynamics of the escape response in bluegill sunfish, *Lepomis macrochirus*. J. Exp. Biol. 211, 3359–3369.

Vaudo, J. J. and Lowe, C. G. (2006). Movement patterns of the round stingray *Urobatis halleri* (Cooper) near a thermal outfall. J. Fish Biol. 68, 1756–1766.

Webb, P. W. (2002). Kinematics of plaice, *Pleuronectes platessa*, and cod, *Gadus morhua*, swimming near the bottom. J. Exp. Biol. 205, 2125–2134.

Wilga, C. D., Maia, A., Nauwelaerts, S. and Lauder, G. V. (2012). Prey handling using whole-body fluid dynamics in batoids. Zoology 115, 47–57.

